# The gut microbiome across the cardiovascular risk spectrum

**DOI:** 10.1101/2023.06.28.546971

**Authors:** Femke M. Prins, Valerie Collij, Hilde E. Groot, Johannes R. Björk, J. Casper Swarte, Sergio Andreu-Sánchez, Bernadien H. Jansen, Jingyuan Fu, Hermie J.M. Harmsen, Alexandra Zhernakova, Erik Lipsic, Pim van der Harst, Rinse K. Weersma, Ranko Gacesa

**Affiliations:** University of Groningen, University Medical Center Groningen, Department of Gastroenterology and Hepatology, Groningen, The Netherlands; University of Groningen, University Medical Center Groningen, Department of Cardiology, Groningen, The Netherlands; University of Groningen, University Medical Center Groningen, Department of Genetics, Groningen, The Netherlands; University of Groningen, University Medical Center Groningen, Department of Pediatrics, Groningen, The Netherlands; University of Groningen, University Medical Center Groningen, Department of Medical Microbiology and Infection prevention, Groningen, The Netherlands; Department of Cardiology, Division of Heart and Lungs, University Medical Center Utrecht, Utrecht, The Netherlands

**Keywords:** gut microbiome, STEMI, cardiovascular risk, coronary artery disease

## Abstract

**Rationale:** Despite significant progress in treatment strategies, cardiovascular disease remains a leading cause of death worldwide. Identifying new potential targets is crucial for enhancing preventive and therapeutic strategies. The gut microbiome has been associated with the development of coronary artery disease (CAD), however our understanding of the precise changes in the gut microbiome occurring during CAD development remains limited.

**Objective:** To investigate microbiome changes in participants without clinically manifest CAD with different cardiovascular risk levels and in patients with ST-elevation myocardial infarction (STEMI).

**Methods and Results:** In this cross-sectional study we characterized the gut microbiome using metagenomics of 411 fecal samples from individuals with low (n=130), intermediate (n=130) and high (n=125) cardiovascular risk based on the Framingham score, and STEMI patients (n=26). We analyzed alpha and beta diversity of the gut microbiome and differential abundance of species and functional pathways among the different groups while accounting for confounders including medication and technical covariates.

Abundances of *Collinsella stercoris, Flavonifractor plautii* and *Ruthenibacterium lactatiformans* showed a positive trend with cardiovascular risk, while S*treptococcus thermophilus* was negatively associated. Furthermore, in the differential abundance analysis we identified eight species and 49 predicted metabolic pathways that were differently abundant among the groups. These species included species linked to inflammation. Starch biosynthesis and phenolic compound degradation pathways were enriched in the gut microbiome of STEMI patients, while pathways associated with vitamin, lipid and amino-acid biosynthesis were depleted.

**Conclusions:** We identified four microbial species that demonstrated a gradual trend in their abundance from low risk individuals to those with STEMI, and species and pathways that were differently abundant in STEMI patients compared to groups without clinically manifest CAD. Further investigation is warranted to gain deeper understanding of their precise role in CAD progression and potential implications, with the ultimate goal of identifying novel therapeutic targets.

**Graphical abstract:** 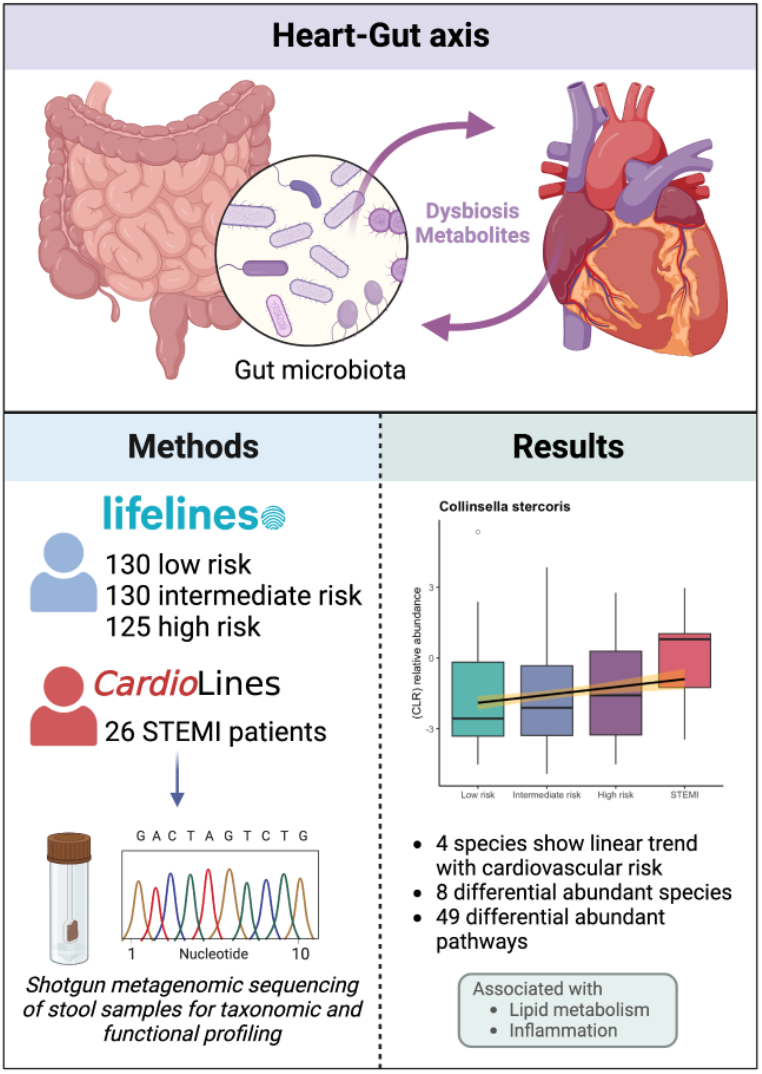

*Created with BioRender.com*

## 1. Introduction

The human gut microbiome, the trillions of microbes that reside in our gastrointestinal tract, plays a vital role in health and disease. This complex system provides us with defense mechanisms against harmful pathogens and aids in nutrient absorption and digestion^1^. In recent years, studies have shed light on the interplay between the gut microbiome and multiple diseases including coronary artery disease (CAD)^2–4^. An imbalance in the microbial communities of the gut microbiome, known as dysbiosis, can trigger systemic inflammation via various mechanisms, and this is believed to play a role in the development of atherosclerosis and pathogenesis of CAD^2,5,6^. In fact, several studies have confirmed the occurrence of dysbiosis in patients with ST-elevation myocardial infarction (STEMI). Specifically, increased abundances of *Proteobacteria, Aerococcacaceae* and *Enterobacteriaceae* have been reported^7,8^. *Enterobacteriaceae* are known to be pro-inflammatory as the interaction of the endotoxin lipopolysaccharide and immune cells triggers an inflammatory response^9^. On the other hand, decreased abundances of *Firmicutes* and *Lactobacillales* have been reported in STEMI patients. These bacteria are recognized for their anti-inflammatory properties because of their involvement in short-chain fatty acids (SCFA) metabolism^7^. Furthermore, it has been observed that STEMI patients show a decreased abundance of the *Lachnospiraceae* family. This beneficial microbe family has been linked to lower serum levels of trimethylamine N-oxide, a gut metabolite recognized to have pro-atherosclerotic effects^8^.

Understanding the complex mechanisms that contribute to disease development and progression is pivotal for developing new preventive and therapeutic strategies for CAD. Therefore, it is important to broaden the scope of investigation beyond conventional cardiovascular risk factors. The potential role of the gut microbiome as a preventive or therapeutic modality for CAD, and as a contributor to cardiovascular risk assessment, is intriguing. However, although dysbiosis signatures have been observed in STEMI patients, there is still a lack of knowledge about the microbial composition in individuals prior to an acute STEMI event. To identify microbial targets for future interventions, it is crucial to determine whether specific changes in microbiome composition occur in the early stages of CAD development.

With this in mind, we studied whether dysbiosis signatures of CAD are already present in individuals at risk for cardiovascular events. Our study aimed to characterize the gut microbiome’s composition across a spectrum of cardiovascular risk profiles, ranging from healthy individuals to those with intermediate/high cardiovascular risk and patients with acute STEMI. We hypothesized that the abundance of microbes reported to be decreased or increased in the STEMI group would demonstrate a gradual trend from low risk individuals to those with STEMI. We analyzed shotgun metagenomes derived from fecal samples of 385 individuals with varying cardiovascular risk profiles and 26 patients with acute STEMI and we compared the metagenomes of different groups on taxonomic and predicted microbial functional levels.

## 2. Methods

### 2.1 Cohort descriptions

For this study we used two Dutch cohorts: (1) myocardial infarction patients participating in the CardioLines Biobank and (2) general population participants of the Dutch Microbiome Project (DMP)^10^. The CardioLines Biobank was established in 2013 and collected body materials from patients with heart disease treated in the University Medical Center Groningen (UMCG, METc number 2012/296). For the present study, inclusion criteria were presence of STEMI, an age older than 18 years, and less than 6 hours since the start of STEMI complaints. Major exclusion criteria were previous myocardial infarction or percutaneous coronary intervention, presence of inflammatory diseases or other heart diseases, usage of anti-inflammatory medication or usage of tavegil/desloratadine. Verbal consent was obtained from all patients at admission, followed by written informed consent. Participants from the DMP were included as groups at risk and a detailed description of the DMP has been published previously^10^. In short, the cohort is part of the Lifelines study (METc number 2017/152), which consists of 167,000 participants from the general population of the northern provinces of the Netherlands for whom stool, blood and metadata information such as diseases and medication are collected. Written consent was signed by all Lifelines participants.

### 2.2. Fecal sample collection

All fecal samples for both cohorts were collected according to the same standardized protocol. The CardioLines samples are samples of the first stool after admission to the hospital for STEMI. Hospital nurses were asked to freeze stool samples directly after production on dry ice and thereafter in the freezer. For the DMP, stool sample collection was performed by participants at home using a standardized stool collection kit provided by the UMCG. The frozen samples were then collected and stored at -80°C until further processing.

### 2.3. Microbial extraction and metagenomic sequencing

Microbial DNA was isolated with the QIAmp Fast DNA Stool Mini Kit (Qiagen, Germany) according to the manufacturer’s instructions, using the QIAcube robot system for automated sample preparation. All CardioLines samples were handled by the same research technicians who handled the DMP samples. Whole-genome shotgun metagenomic sequencing and library preparation was performed at NovoGene (Hong Kong) using the Illumina HiSeq 2000 sequencing platform.

### 2.4. Metagenomic data processing

Metagenomic sequencing reads mapping to Illumina adapters and low-quality reads were trimmed and filtered out using KneadData tools (v0.5.1). The Bowtie2 tool (v2.3.4.1)^11^ was used to remove reads aligning to the human genome (hg19). For the resulting reads, the MetaPhlAn (v3) tool was used to generate taxonomic abundance profiles and the HUMAnN3 pipeline (v0.11.1) for profiling microbial pathway abundances^12^. Taxonomic profiles were subsequently filtered on bacterial and archaeal features. The dataset for the CardioLines cohort contained 526 taxa (2 kingdoms, 10 phyla, 22 classes, 30 orders, 48 families, 107 genera and 256 species). That for the DMP cohort consisted of 942 taxa (2 kingdoms, 12 phyla, 23 classes, 36 orders, 60 families, 137 genera and 406 species).

### 2.5. Group stratification and description

Participants from the DMP were stratified into three groups based on cardiovascular risk as determined by their Framingham score and medical history. Framingham score was calculated using the R package *CVrisk* (v1.1.0)^13^. This score computes the 10-year risk for atherosclerotic cardiovascular disease events using the following parameters: sex, age, total cholesterol, HDL-cholesterol, systolic blood pressure, blood pressure medication, current smoking and diabetes. Risk groups were matched (5:1) with STEMI patients (n=26) based on age, sex, and BMI (factors known to affect the gut microbiome) using the R package *MatchIt* (v4.5.1, method=nearest)^14^, resulting in the following groups:

1. Low risk (n=130): DMP participants with no pre-existing disease, no medication use and a Framingham score <10%.
2. Intermediate risk (n=130): DMP participants without myocardial infarction and with a Framingham score of 10-19%.
3. High risk (n=125): DMP participants without myocardial infarction and with a Framingham score of 20% or higher.

### 2.6. Description of the microbiome

We started our investigation with computation of diversity measures. To achieve this, we merged taxonomy data from both cohorts and the metabolic pathways, keeping the features that were present in both cohorts. Species that did not overlap between cohorts were examined for their mean relative abundance and prevalence, as documented in Supplementary Tables 3 and 4. Of these species, we decided to keep features with a prevalence >5% in the respective cohort (one species) for the calculation of alpha diversity.

For alpha diversity, defined as the diversity of species within a sample (considering its richness, number of different taxonomic groups and evenness and the distribution of abundances of the taxonomic groups), we computed the first three Hill numbers: richness, Shannon diversity index, and the inverse of the Simpson’s index. As one moves from richness to the Simpson’s index, rare species are increasingly penalized. We computed these indices with the *diversity* function in the R *vegan* (v2.5-7)^15^ package. Statistical differences in mean diversities between groups were tested using the non-parametric Wilcoxon-Mann-Whitney test.

Metagenomic sequencing generates compositional data, which means that information can only be acquired as relative abundances, independent of the absolute microbial load in a sample. That means that if one microbial feature increases, other features decrease accordingly^16^. Not taking the nature of this data into account may lead to false positive results. To facilitate statistical testing for the following analyses, we applied the centered log-ratio (CLR) transformation, which put simply, compares the relative abundance of each microbiome in a sample to the “average” microbe in the same sample. To deal with zero values, we implemented a pseudocount by calculating it as half the minimum abundance. Furthermore, we applied a prevalence filter of ≥10%, resulting in 136 species that were kept for subsequent analyses.

Beta diversity, a term describing the extent of difference in microbial composition between samples, was assessed using the Aitchison distance, which corresponds to the Euclidean distance of CLR-transformed relative abundances. For this we used the *vegdist* function in the R *vegan* package. To visualize our data and to examine any potential clustering of the samples, we used principal coordinate analysis (PCoA). PCoA is a dimension-reduction technique used for analysis of high dimensional data. Principal coordinates were constructed and plotted with the *cmdscale* function. The means of the first five principal components were tested using a Wilcoxon test for each pairwise group comparison (Supplementary Table 5). To test the community configurations observed in the PCoA plots, we performed Permutational Multivariate Analysis of Variance (PERMANOVA), using the *adonis2* function from the R *vegan* package with 1000 permutations. We tested the proportion of variance in beta diversity explained by phenotypes in a univariable and a multivariable analysis. In the multivariable analysis, we included covariates previously shown to influence the composition of the microbiome, specifically the use of proton pump inhibitors (PPI) and sequence read depth.

### 2.7. Post-hoc analysis of risk score

Following the above-mentioned analyses, we aimed to investigate the specific effects of the individual components within the Framingham risk score. Here, we analyzed whether certain components of the risk score had a substantial impact on the significant results we observed. To address this, we performed PERMANOVA analyses once more, this time including all eight components of the Framingham score as the variables of interest.

### 2.8. Differential abundance analysis

Differential abundance analysis is a method that can detect variations in the relative abundances of taxonomic groups or pathways between different groups, such as between healthy and diseased individuals. We used linear models to investigate the differential abundance while taking into account the covariates PPI use and sequence read depth (*CLR abundance* (*taxon/pathway*) ∼ *risk_group* + *protin_pump_inhibitor + reads*). To compare groups, we used the *emmeans* (v1.8.6)^17^ package in R, which calculates the estimated marginal means of each group from the fitted model. Any comparison can then be defined, comparing the marginal means between two groups. In total, we performed seven comparisons, including six pairwise comparisons and a ‘merged-group’ comparison that consisted of the low, intermediate and high risk groups grouped together versus the STEMI group. For each contrast, we used the Benjamini-Hochberg false discovery rate (FDR) to correct for multiple testing, with significance set at an FDR<0.05.

## 3. Results

### 3.1 Clinical characteristics

Table 1 presents the clinical characteristics of the study population. In this study we included four distinct groups: 130 individuals from the general population with a low cardiovascular risk (mean age 59.9 years, 77% males), 130 individuals with an intermediate cardiovascular risk (mean age 62.3 years, 100% males), 125 individuals with a high cardiovascular risk (mean age 65.4 years, 100% males) and 26 STEMI patients (mean age 63.5 years, 85% males). More than half of the study participants (56.2%) reported a history of current or former smoking. In terms of medication usage, the intermediate and high risk groups showed some use of medication, especially beta blockers and statins (respectively 23.2% and 19.2% in the high risk group).

**Table 1:** Clinical characteristics of the different groups. Values for categorical variables = counts (proportion, %) for the presence of that phenotype and for numerical variables = mean (SD). CVA = cerebral vascular accident, HDL = high density lipoprotein, LDL = low density lipoprotein, ACE = angiotensin-converting enzyme.

| Phenotype | Low risk group (130) | Intermediate risk group (130) | High risk group (125) | STEMI group (26) | Overall | FDR (BH) - Kruskal Wallis |
| --- | --- | --- | --- | --- | --- | --- |
| Age (years) | 59.9 (7.57) | 62.3 (7.62) | 65.4 (6.29) | 63.5 (11.5) | 62.6 (7.82) | <0.001 |
| Sex (male) | 100 (76.9%) | 130 (100%) | 125 (100%) | 22 (84.6%) | 377 (91.7%) | <0.001 |
| Body Mass Index (kg/m<sup>2</sup>) | 26.8 (3.94) | 26.0 (3.97) | 27.5 (3.42) | 26.7 (3.71) | 26.8 (3.82) | 0.001 |
| Systolic blood pressure (mmHg) | 131 (14.6) | 142 (16.1) | 153 (17.8) | 127 (19.5) | 141 (18.7) | <0.001 |
| Diastolic blood pressure (mmHg) | 75.9 (8.31) | 80.2 (10.2) | 82.7 (9.90) | 81.8 (12.7) | 79.7 (10.1) | <0.001 |
| Heart rate (beats/min) | 66.3 (9.70) | 68.4 (12.2) | 70.8 (13.7) | 78.0 (16.4) | 69.1 (12.6) | 0.002 |
| Smoking | 50 (38.5%) | 73 (56.2%) | 91 (72.8%) | 17 (65.4%) | 231 (56.2%) | <0.001 |
| Diabetes | 0 (0%) | 8 (6.2%) | 34 (27.2%) | 0 (0%) | 42 (10.2%) | <0.001 |
| Hypercholesterolemia | 0 (0%) | 35 (26.9%) | 51 (40.8%) | 4 (15.4%) | 90 (21.9%) | <0.001 |
| Hypertension | 0 (0%) | 39 (30.0%) | 59 (47.2%) | 4 (15.4%) | 102 (24.8%) | <0.001 |
| Artery disease | 0 (0%) | 1 (0.8%) | 4 (3.2%) | 2 (7.7%) | 7 (1.7%) | 0.023 |
| CVA | 0 (0%) | 4 (3.1%) | 3 (2.4%) | 0 (0%) | 7 (1.7%) | 0.240 |
| Leukocytes (10<sup>9</sup>/L) | 5.71 (1.47) | 6.37 (1.71) | 6.98 (1.61) | 11.5 (3.50) | 6.67 (2.23) | <0.001 |
| Creatinine (umol/L) | 87.0 (13.7) | 91.2 (12.6) | 95.1 (17.2) | 77.7 (11.2) | 90.2 (15.1) | <0.001 |
| Total cholesterol (mmol/L) | 5.32 (0.839) | 5.47 (0.975) | 5.31 (1.13) | 5.18 (0.851) | 5.36 (0.980) | 0.325 |
| HDL (mmol/L) | 1.53 (0.376) | 1.24 (0.303) | 1.09 (0.325) | 1.12 (0.320) | 1.28 (0.380) | <0.001 |
| **LDL (mmol/L)** | 3.53 (0.768) | 3.77 (0.864) | 3.55 (1.05) | 3.93 (0.821) | 3.64 (0.902) | 0.033 |
| **Triglyceride (mmol/L)** | 1.11 (0.562) | 1.69 (1.05) | 2.25 (2.20) | 1.19 (0.912) | 1.65 (1.47) | <0.001 |
| **Proton pump inhibitor** | 0 (0%) | 17 (13.1%) | 17 (13.6%) | 2 (7.7%) | 36 (8.8%) | <0.001 |
| **Beta blocker** | 0 (0%) | 11 (8.5%) | 29 (23.2%) | 1 (3.8%) | 41 (10.0%) | <0.001 |
| **Angiotensin antagonist** | 0 (0%) | 8 (6.2%) | 14 (11.2%) | 0 (0%) | 22 (5.4%) | 0.001 |
| **Calcium antagonist** | 0 (0%) | 6 (4.6%) | 10 (8.0%) | 0 (0%) | 16 (3.9%) | 0.103 |
| **ACE inhibitor** | 0 (0%) | 11 (8.5%) | 17 (13.6%) | 2 (7.7%) | 30 (7.3%) | 0.001 |
| **Antithrombotics** | 0 (0%) | 13 (10.0%) | 15 (12.0%) | 0 (0%) | 28 (6.8%) | 0.001 |
| **P2Y12 inhibitors** | 0 (0%) | 1 (0.8%) | 0 (0%) | 0 (0%) | 1 (0.2%) | 0.556 |
| **Diuretics (low ceiling)** | 0 (0%) | 6 (4.6%) | 20 (16.0%) | 0 (0%) | 26 (6.3%) | <0.001 |
| **Antiarrhythmics** | 0 (0%) | 2 (1.5%) | 0 (0%) | 0 (0%) | 2 (0.5%) | 0.250 |
| **Lipid-lowering meds** | 0 (0%) | 0 (0%) | 4 (3.2%) | 0 (0%) | 4 (1.0%) | 0.033 |
| **Statin** | 0 (0%) | 10 (7.7%) | 24 (19.2%) | 0 (0%) | 34 (8.3%) | <0.001 |
| **Oral antidiabetic** | 0 (0%) | 6 (4.6%) | 18 (14.4%) | 0 (0%) | 24 (5.8%) | <0.001 |
| **Insulin** | 0 (0%) | 1 (0.8%) | 5 (4.0%) | 0 (0%) | 6 (1.5%) | 0.047 |
| **Inhalation medication** | 0 (0%) | 1 (0.8%) | 1 (0.8%) | 0 (0%) | 2 (0.5%) | 0.746 |
| **Reads (x10<sup>6</sup>)** | 26.4 (5.6) | 26.8 (3.73) | 26.4 (3.3) | 19.6 (4.05) | 26.1 (4.62) | <0.001 |

### 3.2 Differences in diversity measures observed in the gut microbiome across different cardiovascular risk profiles

We investigated the diversity of the microbiome on species-level to determine whether the microbial community composition and complexity differed among the study groups. We first analyzed the alpha diversity. The measure of richness, which reflects the total number of unique bacterial taxa present in a sample, demonstrated a significant difference between the low risk group and the STEMI group (mean 66.8, SD=11.1 versus mean 72.2, SD=10.5, p=0.034) and the high risk group and the STEMI group (mean 66.2, SD=13.9 and mean 72.2, SD=10.5, p=0.047), also when correcting for sequencing read depth. For the intermediate risk group versus other groups, no significant differences were observed. Our analysis using the Shannon and Simpson’s diversity measures, which not only accounted for taxa richness but also for evenness, did not reveal any differences in alpha diversity between groups, suggesting no distinct microbial compositions. Next, using the inter-sample Aitchison distance (beta diversity), we visualized differences in taxonomic composition through PCoA. No evident clustering of groups was observed in the first five principal components (Figure 1). However, when comparing the centroids of the groups in PCs 2-4, we did observe significant differences, suggesting a relation between inter-individual distance and risk groups (Figure 1).

**Figure 1:**
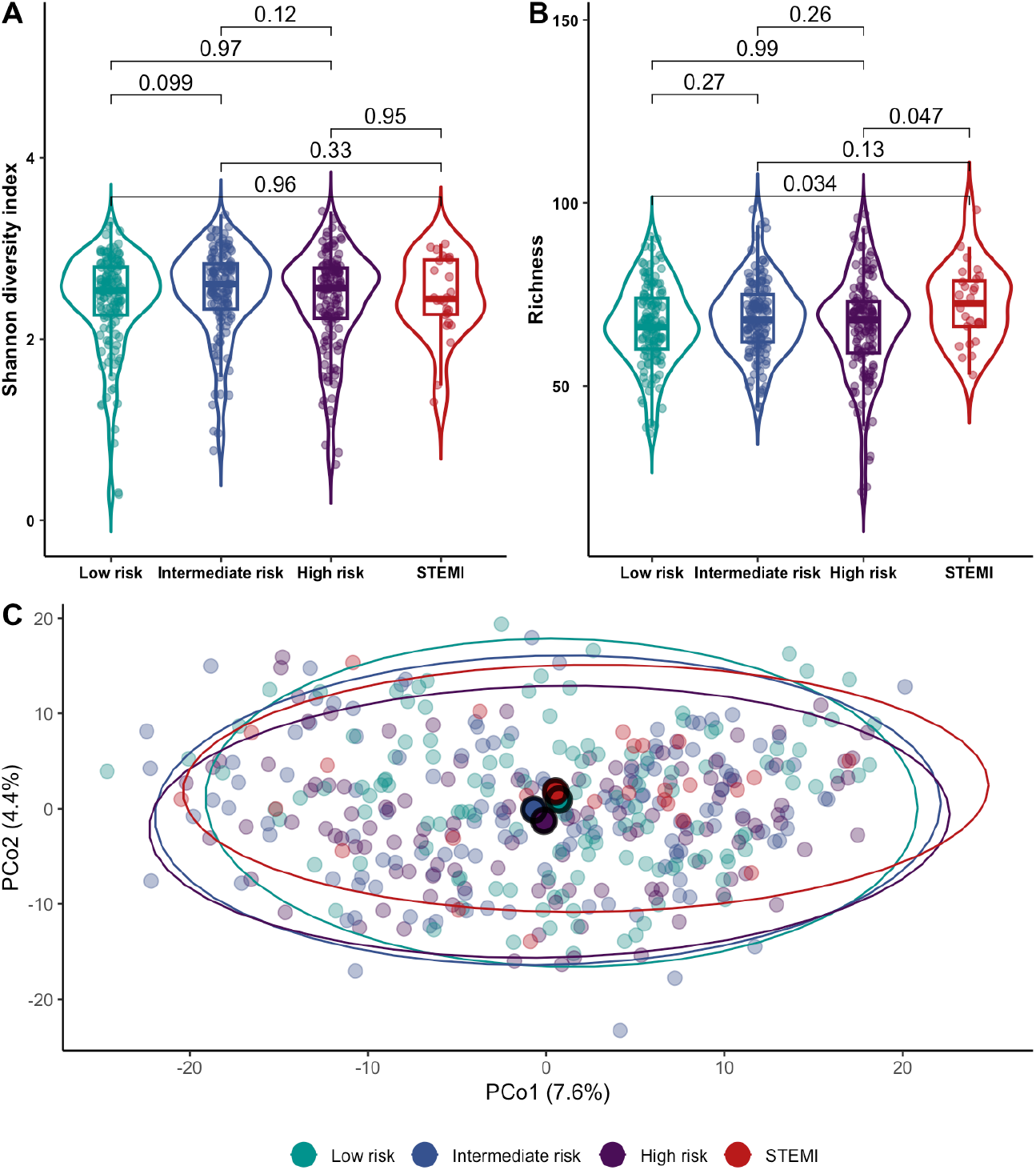
Alpha and beta diversity measures across groups with varying cardiovascular risk. A) Violin plots illustrating the Shannon diversity index for four groups: low risk, intermediate risk, high risk and STEMI. The p-values of the pairwise Wilcoxon tests are shown, which show no significant differences between groups. B) Violin plots that show the richness of each group. The p-values resulting from Wilcoxon tests for each comparison are indicated. Low risk versus STEMI and high risk versus STEMI are significantly different (p = 0.034 and p = 0.047) C) Principal coordinates plot depicting the Aitchison distances at species level, using CLR-transformed relative abundance data. Individual data points are represented by green dots for low risk individuals, blue dots for intermediate risk individuals, purple dots for high risk individuals and red dots for STEMI patients. The large circle represents the centroid, which represents the mean composition of each group.

### 3.3 Cardiovascular risk profiles, based on the Framingham score, explain part of the variance in the gut microbiome

We subsequently conducted PERMANOVA analyses to test the proportion of variance of the gut microbiome that could be explained by the cardiovascular risk profile. In a multivariable analysis, in which we included the covariates PPI use and sequence read depth, risk profile explained a significant proportion of the microbiome inter-sample distance variance (R2=0.011, p≤0.001). This indicates that there are significant differences in microbial composition between groups. As a post-hoc analysis, we evaluated the individual variables of the Framingham risk score to determine if the results we observed were driven by a specific variable. Here we found that four out of eight risk-score variables significantly explained variance in taxonomic composition; diabetes (R2=0.007, p≤0.001), age (R2=0.006, p≤0.001), smoking (R2=0.004, p=0.022) and HDL (R2=0.003, p=0.024).

### 3.4 Decreased abundance of anti-inflammatory species and increased abundance of lipid-metabolism-associated species in the STEMI group compared to groups with lower cardiovascular risk

We used linear models to investigate trends in the relative abundances of species across cardiovascular risk profiles with increasing risk. Here we found three species that showed a positive association with increasing cardiovascular risk (*Collinsella stercoris*, coeff=0.27, FDR=0.049; *Flavonifractor plautii*, coeff=0.53, FDR=0.039 and *Ruthenibacterium lactatiformans*, coeff=0.39, FDR=0.003), while *Streptococcus thermophilius* showed a negative association (coeff=-0.33, FDR=0.0499).

We then performed a differential abundance analysis using linear models in which we tested six pairwise comparisons between all groups and one comparison of the low, intermediate and high risk groups grouped together versus the STEMI group. While our previous analysis tested for taxa that linearly increased or decreased their abundance between groups, this analysis allows for non-linear changes between groups. In total, we found eight unique species showing differential abundance between cardiovascular risk groups (FDR<0.05, Figure 2). Among these species, certain ones showed differential abundance in multiple comparisons, and two species were overlapping with those detected in the linear analysis. We found two species to be differently abundant in the low risk versus STEMI comparison (*R. lactatiformans* FDR=0.029 and *C. stercoris* FDR=0.008), six species when comparing intermediate risk vs STEMI (*Rothia mucilaginosa* FDR=0.049, *Odoribacter splanchnicus* FDR=0.049, *C. stercoris* FDR=0.005, *Bacteroides vulgatus* FDR=0.008, *Bacteroides uniformis* FDR=0.008 and *Agathobaculum butyriciproducens* FDR=0.033), four species when comparing high risk versus STEMI (*Ruminococcaceae bacterium D5* FDR=0.044, *C. stercoris* FDR=0.019, *B. vulgatus* FDR=0.014 and *B. uniformis* FDR=0.019) and four species were differently abundant when comparing STEMI vs all other groups. Focusing on the species that were differently abundant in the STEMI group compared to all other groups, we observed *C. stercoris*, a species that has been associated with cholesterol metabolism (displaying a negative correlation with serum HDL-cholesterol and a positive correlation with triglycerides^18^), to be enriched in the STEMI group (FDR=0.005). *A. butyriciproducens*, a butyric acid-producing bacteria, was also enriched in the STEMI group compared to other groups (FDR=0.037). A lower relative abundance of *Bacteroides* species (*B. vulgatus* FDR=0.013, *B. uniformis* FDR=0.019), known for their ability to produce SCFAs, was observed in the STEMI group compared to all other groups.

**Figure 2:**
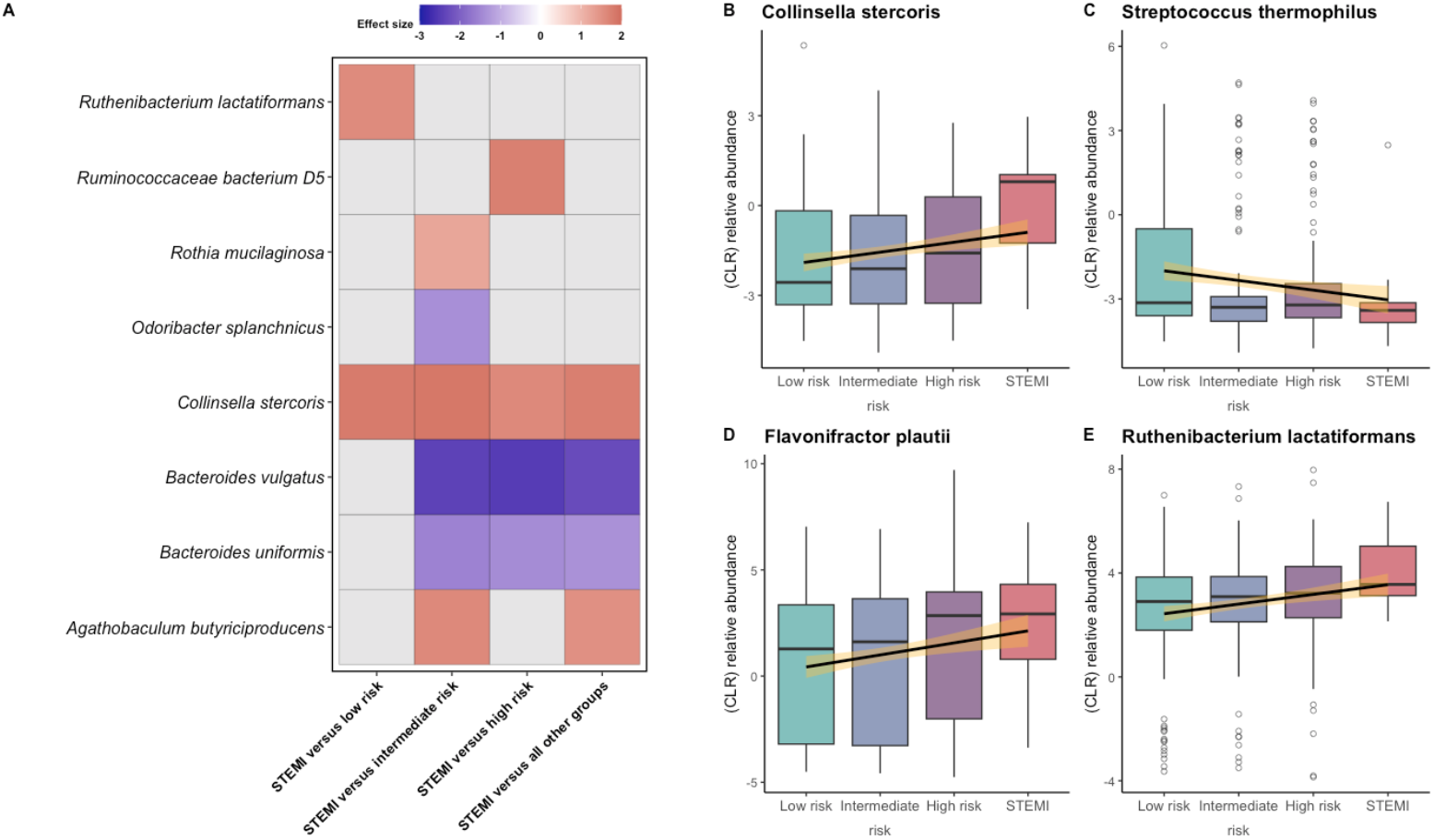
Differential abundant species. A) Heatmap representing significant findings from pairwise differential abundance analyses using linear models. The red color indicates higher abundance of the species in the STEMI group, while blue indicates a lower abundance in the STEMI group. B,C,D,E) CLR-transformed relative abundance of the four species that show a significant linear trend in their abundance across the different groups. The black line represents the trend line, and the yellow shadow indicates the standard error. The boxplots show the relative abundance in each group.

### 3.5 Multiple pathways related to vitamin biosynthesis and amino acid biosynthesis are decreased in the STEMI group

To examine the functional potential of the microbial community in individuals with varying levels of cardiovascular risk, we analyzed the relative abundance of metabolic pathways. We selected pathways that were present in ≥10% of the samples and used a linear model for pairwise comparisons between groups. Of the 408 pathways identified, 49 unique pathways had distinct differences in abundance between the different cardiovascular risk groups (Figure 3). Of these, only four pathways were increased in the STEMI group in comparison to groups with lower risk, while the majority of the pathways were decreased in the STEMI group in contrast to other groups. These decreased pathways were associated with many processes such as nucleotide biosynthesis, fatty acid and lipid biosynthesis and carbohydrates biosynthesis. We observed a decrease in four amino acid biosynthesis pathways in the STEMI group. These pathways are involved in: 1) L-citrulline biosynthesis, linked to nitric oxide synthesis and suggested to reduce blood pressure through supplementation^19^, 2) aspartate/asparagine biosynthesis, shown to promote production of inflammatory cytokines^20^, 3) L-orthinine biosynthesis, linked to a higher incidence of cardiovascular death when the L-arginine/L-orthinine ratio is low^21^ and 4) L-lysine biosynthesis, where a previous study found that individuals with pre-existing type 2 diabetes and elevated lysine levels are at the highest risk of developing cardiovascular disease^22^. In addition, we found six vitamin biosynthesis pathways decreased in the STEMI group including vitamin B7 (biotin), vitamin B1 derivatives and vitamin B6.

**Figure 3:**
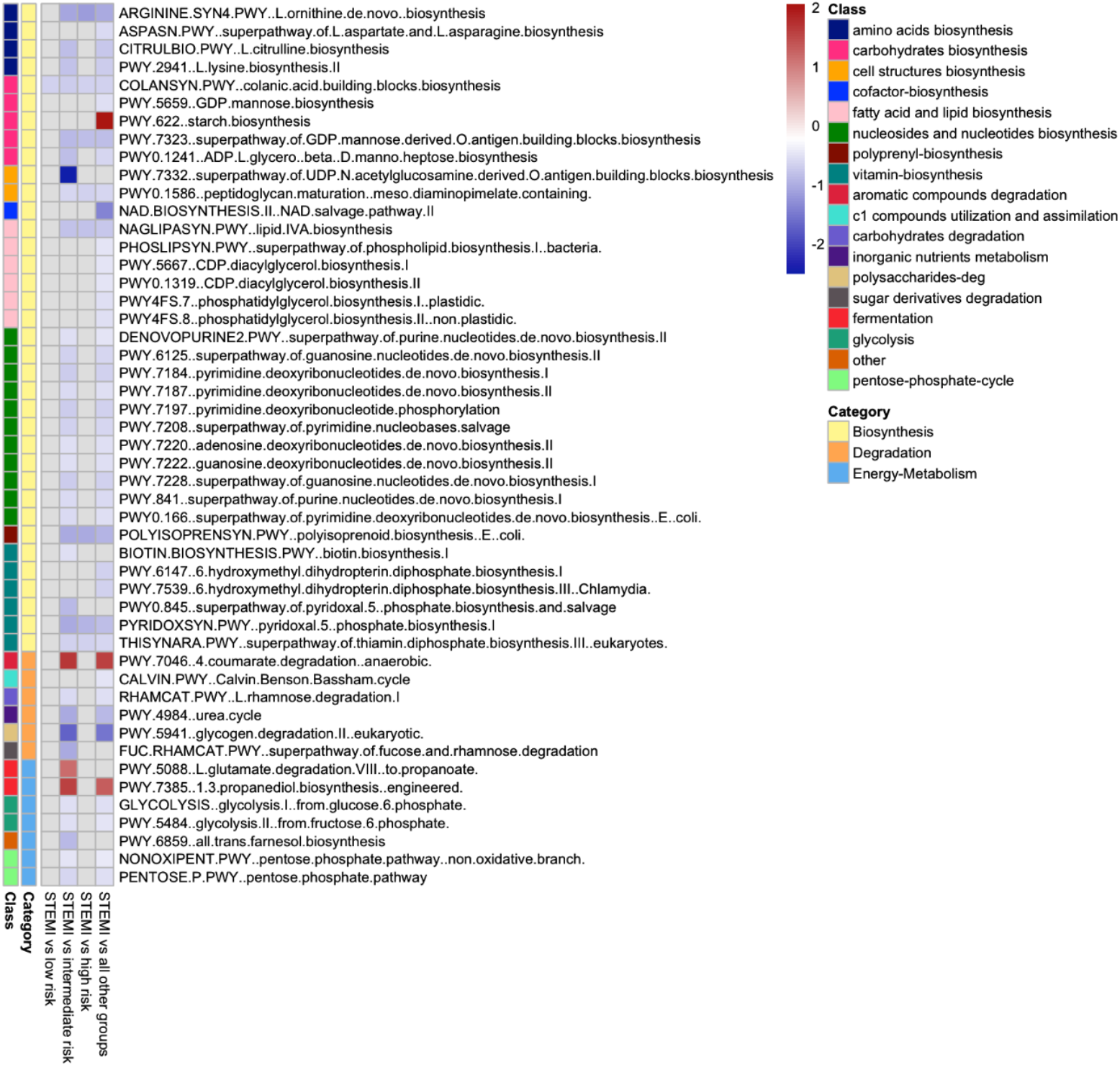
Differential abundant microbial predicted pathways across groups. Linear models analysis identified 49 predicted microbial pathways that showed significant alterations in relative abundance between groups (FDR < 0.05). The heatmap displays the effect size, where red indicates an increased abundance in the STEMI group, while blue indicates a decreased abundance. Additionally, the left-side color-coded columns show the functional classes and categories to which each pathway belongs.

## 4. Discussion

In this study we explored the microbial composition and functional potential of the gut microbiome in groups with varying degrees of cardiovascular risk and STEMI patients. Our analysis revealed four species that showed a significant linear trend in their abundance from low risk individuals to STEMI patients. *C. stercoris, F. plautii* and *R. lactatiformans* showed an increasing abundance with higher cardiovascular risk, whereas *S. thermophilius* showed a decreasing abundance. Furthermore, we found eight species and 49 pathways to be differently abundant among the groups. Regarding alpha diversity measures, we observed differences in richness between the STEMI group and both the low and high risk groups. Additionally, the risk groups accounted for a significant proportion of the microbiome inter-sample distance variance.

An intriguing finding is the gradual increase in the abundance of the species *C. stercoris* in groups with increasing cardiovascular risk. As previously reported, patients with high cardiovascular risk show an increased abundance of the genus *Collinsella*^23^. These microbes are believed to modulate host metabolism by affecting intestinal cholesterol absorption, decreasing glycogenesis in the liver and increasing triglyceride synthesis^5^. The role of *Collinsella* in cholesterol metabolism was confirmed by a study that demonstrated a positive correlation between serum cholesterol and *Collinsella*^24^. Given that hypercholesterolemia is a well-established risk factor for CAD, an increased abundance of *Collinsella* could contribute to the risk of developing CAD by promoting hypercholesterolemia. Nevertheless, not all reports on *Collinsella* in CAD present consistent findings; a reduction in the abundance of *Collinsella* in coronary heart disease patients has also been reported, but this was suggested to be due to treatment with statins^25^. In our study, some participants received statin treatment, however a previous study found little effect of statins on the overall composition of the microbiome, and these effects were not specifically associated with *Collinsella*^26^.

Bacterial species that were significantly decreased in the STEMI group included *B. vulgatus, B. uniformis* and *O. splanchnicus*. These findings are consistent with previous studies that reported decreased *Bacteroides* abundance in patients with CAD^25^. *Bacteroides* are common microbes in the gut and are considered commensal bacteria. It has been suggested that *B. vulgatus* has an anti-inflammatory effect by affecting microbial lipopolysaccharide synthesis^27^. *B. uniformis* has been studied in a mouse model, where its administration was found to reduce serum cholesterol and triglyceride levels, improve glucose metabolism and insulin sensitivity, reduce leptin levels and enhance immune function^28^. *O. splanchnicus* is a common SCFA producer, and a reduced species abundance has been associated with diseases such as metabolic syndrome^29^. Higher abundance of the genus *Odoribacter* has been linked to lower blood pressure in overweight and obese pregnant women^30^.

In addition, in STEMI patients, we observed an enrichment of species belonging to the *Oscillospiraceae* family (previously names *Ruminococcacaea*), which includes bacteria known for their butyrate production. To the best of our knowledge, the three species identified in this study that belong to this family (*R. lactatiformans, A. butyriciproducens* and *Ruminococcaceae bacterium D5)* have not previously been described in relation to cardiovascular disease, although other members of this family have been reported in that context. Previous studies have reported contradictory findings regarding the abundance of the *Oscillospiraceae* family. Some studies found increased abundance of species of this family in patients with severe (cardiac) disease, while others noted decreased abundance in CAD patients^25,31,32^. This discrepancy could reflect individual species within the same family having distinct functions. Additionally, a novel finding of our study is the increased abundance in STEMI patients of *Rothia mucilaginosa*, a pathogenic species typically found in the oral cavity and has been associated with many infections such as meningitis and endocarditis^33^. To increase our understanding of the differentially abundant species identified in this study, it is crucial that future studies are conducted to replicate these findings.

We used metagenomic data to examine the functional potential of the microbiome in each group. We found that in the STEMI group, compared to the other risk groups, microbial pathways related to carbohydrate biosynthesis, fermentation and energy metabolism were enriched. Of the pathways decreased in the STEMI group, many were related to nucleotides, vitamin, amino acids and carbohydrate biosynthesis. One pathway of interest is the *pyridoxal 5’-phosphate biosynthesis I* pathway, which is involved in the biosynthesis of vitamin B6. Vitamin B6 is an essential micronutrient known for its anti-inflammatory properties and is involved in more than 160 different biochemical processes. Previous studies have reported significant associations between low vitamin B6 levels and increased risk of coronary heart disease. More specifically, a significant inverse linear relation of plasma pyridoxal 5’-phosphate (the active coenzyme form of vitamin B6) levels and risk of coronary heart disease in women has been reported^34^. However, it should be emphasized that the decreased abundance of this pathway in the STEMI group is for the microbial metabolic pathway not the human pathway. We did not assess vitamin B6 or pyridoxal 5’-phosphate levels in the current study. As a result, it remains unknown if the observed microbial pathway abundance correlates with reduced serum levels in the host.

This study has several strengths. Firstly, we used shot-gun sequencing to obtain metagenomic data, which provides improved taxonomic resolution compared to the more commonly used 16S rRNA gene sequencing techniques. Furthermore, this approach allowed us to investigate functional profiles of metabolic pathways of the microbial communities. Secondly, the inclusion of participants from a general population cohort for the risk groups ensures that selection bias is minimized. In addition, the availability of comprehensive metadata from this cohort made it possible to calculate risk scores. Thirdly, a notable strength is that the STEMI group had minimal medication usage, as cardiovascular medication was only started after the acute event and sample collection. This represents a common scenario in which patients presenting with STEMI are often not previously known by a cardiologist, resulting in the absence of medication regimens before the event.

However, our study also has some limitations that should be addressed. Firstly, our STEMI group had a relatively small sample size of 26 patients. Future studies would benefit from a larger sample size for the groups to increase statistical power. Secondly, the intermediate and high risk groups included in the study consisted only of male participants. In future studies an alternative cohort including women with cardiovascular risk should be considered to make the findings more generalizable. Thirdly, the complex and multifactorial nature of the crosstalk between the gut microbiome and host in CAD makes it difficult to determine causality when using *in silico* analyses alone. Therefore, future studies should include experimental designs, such as randomized trials with interventions and culturing of fecal bacteria to provide further insights into the directional and causal relationships of the gut microbiome and CAD. Lastly, we are aware that diet and lifestyle can significantly influence the gut microbiome, but the present study lacked data on these factors and we could not adjust for them.

Overall, we explored if dysbiosis signatures of the gut microbiome seen in CAD/STEMI patients could be detected in individuals at risk. We identified four species to show a gradual trend in abundance with increasing cardiovascular risk and eight species to be differently abundant in the STEMI group compared to individuals at risk. These results contribute to the existing body of research on the relationship between CAD and the gut microbiome and offer insight into potential therapeutic directions for future mechanistic and interventional studies.

## Acknowledgments

The authors wish to acknowledge the services of the Lifelines Cohort Study, the contributing research centers delivering data to Lifelines, and all the study participants. We would also like to thank all STEMI patients from the Cardiolines cohort for fecal sample collection. We would also like to express our appreciation to Kate McIntyre (Scientific Editor, Departments of Genetics, University Medical Center Groningen) for her valuable English editing assistance.

## Sources of Funding

The Lifelines Biobank initiative was made possible by a subsidy from the Dutch Ministry of Health, Welfare and Sport; the Dutch Ministry of Economic Affairs; the University Medical Center Groningen (UMCG), the Netherlands; the University of Groningen and the Northern Provinces of the Netherlands. Some of the authors of this project received support from various funding agencies; however, this funding was not relevant to the content of this manuscript. J.F. is supported by the Dutch Heart Foundation IN-CONTROL (CVON2018-27), the ERC Consolidator grant (grant agreement No. 101001678), NWO-VICI grant VI.C.202.022, and the Netherlands Organ-on-Chip Initiative, an NWO Gravitation project (024.003.001) funded by the Ministry of Education, Culture and Science of the government of The Netherlands. A.Z. is supported by the Dutch Heart Foundation IN-CONTROL (CVON2018-27), the ERC Starting Grant 715772, NWO-VIDI grant 016.178.056, and the NWO Gravitation grant Exposome-NL (024.004.017). R.K.W. is supported by the Seerave Foundation, the Netherlands Organization for Scientific Research (NWO), and the EU Horizon Europe Program grant miGut-Health: personalized blueprint of intestinal health (101095470). The funders had no involvement in the study design, data collection and analysis, decision to publish, or preparation of this manuscript

## Disclosures

R.K.W. acted as consultant for Takeda, received unrestricted research grants from Takeda, Johnson & Johnson, Tramedico, and Ferring, and received speaker fees from MSD, Abbvie, and Janssen Pharmaceuticals. All other authors declare that they have no relevant competing financial interests or relationships that could have influenced the work reported in this paper.

## Novelty and Significance

What is known?

- The gut microbiome is associated with the development of atherosclerosis and pathogenesis of coronary artery disease (CAD)
- Dysbiosis is present in the gut microbiome of ST-elevation myocardial infarction (STEMI)

What new information does this article contribute?

- The findings show varying gut microbial species abundance between STEMI patients and groups at cardiovascular risk
- *Collinsella stercoris, Flavonifractor plautii* and *Ruthenibacterium lactatiformans* are increasingly abundant, while *Streptococcus thermophilus* is decreasingly abundant across the cardiovascular risk profiles.
- An increased abundance of *Ruthenibacterium lactatiformans, Agathobaculum butyriciproducens, Ruminococcaceae bacterium D5* and *Rothia mucilaginosa* in STEMI patients compared to groups at risk has not been reported before.

Despite previous studies demonstrating dysbiosis in STEMI patients, our understanding of the precise microbiome changes across the cardiovascular risk spectrum remains limited. This study addresses this knowledge gap by providing insights into the gut microbiome composition of individuals across varying cardiovascular risk levels and STEMI patients. By examining the gut microbiome of carefully selected participants from the general population with three different risk levels and a unique group of STEMI patients, we identified microbial species and pathways with differential abundance across the groups. Several of these species and pathways are associated with inflammation and lipid metabolism, which are key factors in CAD development. Our findings did not only contribute to the growing body of evidence on the involvement of the gut microbiome in the cardiovascular risk spectrum, but also presents promising prospects for future advancements. Once a causal association has been established, the ultimate goal would be to use these differentially abundant species for the development of personalized risk assessment tools and novel non-invasive microbiome based interventions for enhancing preventative and therapeutic strategies for CAD in the future.

## Notes

### Competing Interest Statement

The authors have declared no competing interest.

